# Molecular evolution of the Angiotensin II receptors AT1 and AT2: Specificity of the sodium binding site in amniota

**DOI:** 10.1101/2021.06.04.446765

**Authors:** Asma Tiss, Rym Ben Boubaker, Daniel Henrion, Hajer Guissouma, Marie Chabbert

## Abstract

In vertebrates, the octopeptide angiotensin II (AngII) is an important *in vivo* regulator of the cardiovascular system. It acts mainly through two G protein-coupled receptors, AT1 and AT2. To better understand the interplay between these receptors throughout the evolution of the renin-angiotensin system (RAS), we combined a phylogenetic study to electrostatics computations and molecular dynamics (MD) simulations of AT1 and AT2 receptors from different species. The phylogenetic analysis reveals a mirror evolution of AT1 and AT2 that are both split in two clades, separating fish from terrestrian receptors. It also indicates that the unusual allosteric sodium binding site of human AT1 is specific of amniota. Other AT1 and AT2 receptors display a canonical sodium binding site with a serine at position 7.46 (Ballesteros numbering). Electrostatics computations and MD simulations support maintained sodium binding to human AT1 with ingress from the extracellular side. Comparison of the sodium binding modes in AT1 and AT2 from humans and eels indicates that the allosteric control by sodium in both AT1 and AT2 evolved during the transition from an aqueous to a terrestrial environment. The unusual S7.46N mutation in amniota AT1 is mirrored by a L3.36M mutation in amniota AT2. The S7.46N mutation increases the specificity of AT1 for AngII relative to Ang derivatives, whereas the L3.36M mutation might have the opposite effect on AT2. Both mutations should contribute to the split of the renin-angiotensin system into the classical (AngII/AT1) and counter-regulatory (Ang1-7/AT2, Mas) arms in amniota.

**AUTHOR SUMMARY:** The analysis of protein sequences from different species can reveal interesting trends in the structural and functional evolution of a protein family. Here, we analyze the evolution of two G protein-coupled receptors, AT1 and AT2, which bind the angiotensin II peptide and are important regulators of the cardiovascular system. We show that these receptors underwent a mirror evolution and that specific mutations of the sodium binding pocket in both AT1 and AT2 occurred in amniota. We underwent electrostatics computations and molecular dynamics simulations to decipher the details of the sodium binding mode in eel and human receptors, as prototype of fish and amniota receptors. Our results suggest that evolution favored an increased specificity of AT1 and a decreased specificity of AT2 for angiotensin II as compared to its derivatives. In turn, these data suggest that mutations in the sodium binding pocket of G protein-coupled receptors might be an efficient way to gain functional evolution.

## INTRODUCTION

The renin-angiotensin system (RAS) is an important *in vivo* regulator of multiple cardiovascular and renal functions in vertebrates [1-3]. It initiates with the cleavage of the angiotensinogen by renin into the decapeptide angiotensin I that is then cleaved by the angiotensin converting enzyme (ACE1) into the octapeptide angiotensin II (AngII). AngII is subsequently cleaved into angiotensin 1-7 (Ang1-7) by the angiotensin converting enzyme (ACE2). A variety of other angiotensin derivatives are produced by diverse enzymes. The RAS system is composed of two axes. In the “classical” axis, AngII binds the type 1 AngII receptor (AT1) to induce most known effects of AngII, including vasoconstriction and increased blood pressure, anti-natriuresis, hypertrophy, fibrosis and inflammation [4,5]. Dysregulation of this axis promotes cardiac and vascular damages and AT1 blockers are widely used to fight hypertension. The “counter-regulatory” axis involves the AngII type 2 receptor (AT2) and the MAS receptor, through AngII (AT2) and Ang1-7 (AT2, MAS). These receptors promote vasodilation, natriuresis, anti-inflammation, anti-fibrosis and anti-proliferative responses and counterbalance AT1 effects [1,6,7].

AT1 and AT2 are two G protein-coupled receptors that share 30% sequence identity and similar affinity for AngII but have very different properties, expression profiles and functions [4]. AT1 is expressed in all organs. When it is stimulated by AngII, it activates both G proteins and β-arrestins. A variety of AT1 ligands induce biased signaling with preferential activation of G proteins or β-arrestins [8,9]. By contrast, AT2 is widely expressed during development, whereas, in the adult life, it is expressed only in a few organs except in pathological conditions where its expression is up-regulated [5,10,11]. Direct AT2 activation of the conventional GPCR effectors (G proteins and β-arrestins) remains controversial and the primary signalling pathways involved in AT2 signaling are unknown [12-14]. A better knowledge of these pathways is important as stimulation of AT2 in pathological situations might be an interesting alternative to the blockade of AT1 [11].

Several structures of AT1 [15-17] and AT2 [18-20] complexed to different ligands have been resolved. Comparison of the structures of AT1 complexed to AngII and biased agonists provide clues to the biased responses of AT1 [16]. AT2 has been crystallized in active states only [18-20], which supports the hypothesis of a “relaxed”, prone to activation conformation [21].

Sodium binding is an essential element of receptor activation of most class A GPCRs. It acts as a negative allosteric regulator, by stabilizing the inactive structure, and might be involved in the activation mechanism [22,23]. Sodium coordination involves a strictly conserved aspartic acid at positions 2.50 (Ballesteros’ numbering [24]) and polar residues at positions 3.35, 3.39, 7.45 or 7.49. AT1 propensity for biased signaling is related to the rotameric orientation of two residues in the sodium binding site that act as activation switch, N3.35 and N7.46 [16]. Asn at position 3.35 is frequent in GPCRs [25]. Asn at position 7.46 is unusual and might prevent sodium binding while stabilizing the inactive structure [16]. By contrast, human AT2 (AT2h) has a conventional sodium binding site, with N3.35 and S7.46 patterns.

Since residues from the sodium pocket have a functionally important role in AT1 activation, a better knowledge of the history of this site is mandatory to understand the role of this unusual sodium pocket not only in the activation mechanism of AT1 but also in the balance between AT1 and AT2 activation. This prompted us to analyze the evolutionary information contained in the extend sequences of both AT1 and AT2.

The evolutionary approach revealed a mirror evolution of AT1 and AT2. Both AT1 and AT2 are split in two clades that separate fish from terrestrian receptors. Subsequent mutations of residues in or close to the sodium binding site occurred specifically in amniota receptors. To understand the effects of these changes, we carried out molecular dynamics simulations to determine (1) the sodium ingress pathway to the canonical binding site in AT1h and (2) the characteristics of the sodium binding site in AT1 and AT2 receptors from humans and eels. The mutations specific of amniota AT1 and AT2 receptors have physiological consequences for the regulation of the renin-angiotensin system that will be discussed.

## RESULTS

### 1. Evolution of the angiotensin receptors

Angiotensin receptors from Uniprot present strong sequence biases with about 30% of all pairs having a sequence identity greater that 80%. It was thus necessary to remove highly similar sequences from the data set to avoid redundancy. Using the NRDB program [26], we clusterized the sequences with a threshold of 90% sequence identity to obtain a non-redundant set. This led to a collapse in the number of sequences (Table I). The 223 AT1 receptors sequences in Uniprot were grouped in 38 clusters from which a single representative sequence was selected. The 120 mammalian AT1 sequences clustered in 3 groups. The cluster containing human AT1 contained a total of 115 sequences. For AT2, the number of sequences decreased from 115 in Uniprot to 37 non-redundant sequences. The 68 mammalian AT2 formed 5 clusters. 38 sequences clustered with human AT2. This classification leads to similar number of non-redundant sequences for AT1 and AT2. Using the non-redundant representatives, 51% and 74% of, respectively, AT1 and AT2 pairs have less than 50% identity (Fig. 1a).

**TABLE I.** Sets of AT1 and AT2 sequences used for the phylogenetic analysis.

|  | Sequences in Uniprot | Clusters with id >90 |
| --- | --- | --- |
| Total AT1 | 223 | 38 |
| Mammalian AT1 | 120 | 3 |
| Cluster including human AT1 | 115 | 1 |
| AT2 | 115 | 37 |
| Mammalian AT2 | 68 | 5 |
| Cluster including human AT2 | 39 | 1 |

**Fig 1:**
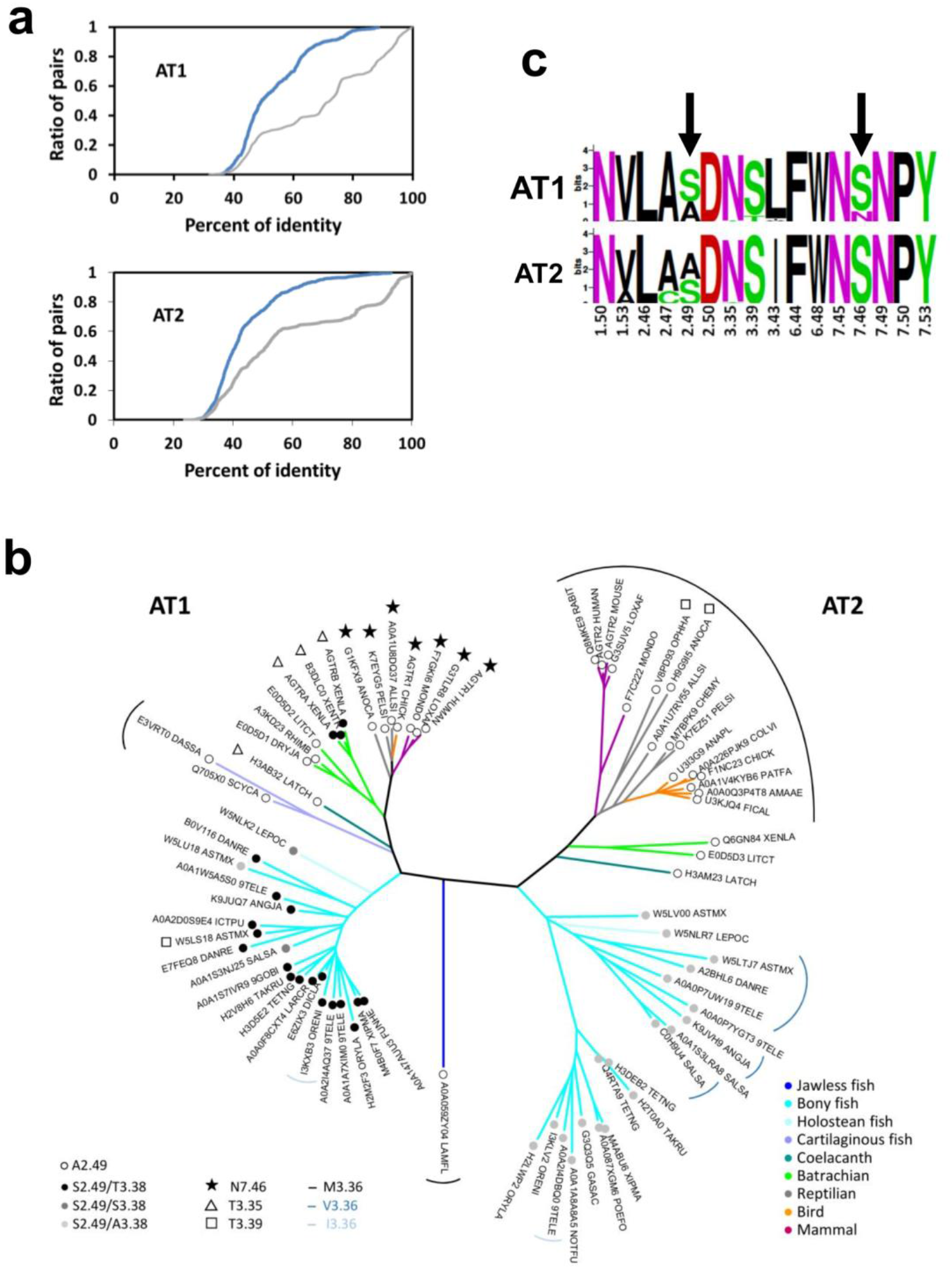
Evolution of the angiotensin receptors AT1 and AT2. (a) Cumulative distribution functions of the pairwise identities for AT1 (top) and AT2 (bottom), in the exhaustive sets obtained from Uniprot (grey lines) and in the non-redundant sets (blue lines); (b) Neighbor-joining tree of the non-redundant set of AT1 and AT2 receptors (500 bootstraps). The color code of the tree branches refers to the phylum (jawless fish: dark blue, bony fish: cyan, holostean fish: light cyan, cartilaginous fish: slate, coelacanth: deepteal, batrachian: green, reptilian: grey, bird: orange, mammal: purple). The external labels indicate mutations in the sodium binding site as compared to the canonical sodium binding site characterized by N3.35, S3.39 and S7.46 (N7.46: stars, T3.35: open triangles, T3.39: open squares). The internal labels indicate the residues at positions 2.49 and 3.38 (A2.49: open circle, S2.49/T3.38: black circle, S2.49/S3.38: dark grey circle, S2.49/A3.38: light grey circle). The circular arcs indicate the residue at position 3.36 (black: M3.36, dark blue: V3.36, light blue: I3.36, no arc: L3.36); (c) Logo plots of the sodium binding sites in AT1 and AT2. For these plots, all the residues lining up the sodium binding site were taken into account [23]. The arrows highlight the mutations at positions 2.49 and 7.46.

Using the aligned sequences of the non-redundant set, we built a Neighbour Joining phylogenic tree of the angiotensin receptors. Starting from a single angiotensin receptor more closely related to AT1 in jawless fish (lamprey), duplication in an ancestor of jawed fishes led to an almost symmetrical evolution of the AT1 and AT2 receptors, hinting at their complementary role in RAS control (Fig. 1b). Each AT1 and AT2 sub-tree possesses two major clades: one for fishes and one for terrestrian animals. There are two exceptions to this pattern: (1) the coelacanth sequences are clearly part of the “terrestrian” clades both for AT1 and AT2, and close to the batracian sequences. This position is consistent with its status of “living fossil” with an intermediary position in the tree of life; (2) only AT1 sequences are present for cartilaginous fishes (catfish, whales) (see Materials et Methods) and are closer to terrestrian than to fish sequences. Interestingly, this feature was observed for the full genome of whales [27].

We analyzed the residues lining the canonical sodium binding site [23] in this non-redundant sequence set. Most residues are highly conserved between and within the AT1 and AT2 sets (Fig. 1c). In particular, residues usually involved in sodium coordination (D2.50, N3.35, S3.39 and N7.45) are highly conserved. A marked difference is observed at position 3.43 with Leu and Ile for AT1 and AT2, respectively. In both sets, the position 2.49 differentiates fishes (S2.49) from terrestrian (A2.49) receptors (Fig. 1b). The C2.47 pattern is present only in some AT2 sequences from fishes. Noteworthy, as observed in most GPCRs, a serine residue at position 7.46 is present in most AT1 and AT2 receptors (Fig. 1b). An Asn at this position is observed only in amniota (reptilians, birds and mammals). This indicates that this specificity of AT1 has been acquired during the transition from an aqueous to a terrestrian environment.

A “canonical” sodium binding site with two Asn residues at positions 3.35 and 7.45 and two Ser residues at positions 3.39 and 7.46 is thus observed in AT2 receptors and in AT1 receptors from fishes and some amphibians (Neobatracia clade). The S7.46N mutation is observed specifically in amniota while other coordinating residues are maintained. Notably, a N3.35T mutation of the residue facing S7.46 is observed in the coelacanth and batracians from the pipoidea clade (Xenopus), suggesting that evolution might have led to different solutions for a similar problem.

### 2. Properties of D2.50 in AT1 and AT2

The deprotonation of D2.50 is a key element of sodium binding to GPCRs. We thus wondered whether the mutations in the sodium binding site of AT1 and AT2 can alter the pKa of D2.50. To answer this question, we selected AT1 and AT2 from *Anguilla japonica* (eel) as prototypes of the fish receptors. We modeled these receptors from crystallographic structures of human AT1 and AT2 (see Materials and Methods) and we computed the pKa of D2.50 in AT1 and AT2 from humans (AT1h, AT2h) and eels (AT1e, AT2e) along with the human AT1 receptor with the N7.46S mutation (AT1m). The pKa of D2.50 in the opioid δ receptor (OPRD) and in the chemokine receptor CXCR4 were also computed as controls. To calculate pKa, we used the DelphiPKa program with the model of Gaussian representation of atomic densities proposed by Alexov and coworkers [28-30]. This representation yields a smooth Gaussian-based dielectric function without defining a molecular surface. Electrostatic computations depend on two parameters: the width of the Gaussian function, σ, and the protein dielectric value, ε_in_. We compared a wide range of parameters (σ varying from 0.5 to 1, ε_in_ varying from 2 to 10) for AT1 and OPRD in search of sets compatible with sodium binding to OPRD (see Materials and Methods). Each pair of parameters led to similar pKa values for AT1 and OPRD (Supplementary Fig. S1), indicating that D2.50 has the same protonation state in both receptors. Sodium binding to OPRD is not compatible with the default values for internal groups (σ = 4, ε_in_ = 0.9) which lead to neutral pKa (7.3 ± 1.0) but supports the default values for external groups (σ = 0.7 and ε_in_ = 8), which lead to a pKa of 4.1 ± 0. These latter parameters are consistent with the location of D2.50 facing a water filled cavity. Using these parameters, we obtained similar values for the pKa of D2.50 in AT1 and AT2 from both humans and eels and in the N7.46S AT1 mutant (Fig. 2a). This indicates that the presence of N7.46 in the sodium binding site does not alter the physico-chemical properties of D2.50 in human AT1. Notably, we observed a significant decrease in the pKa of D2.50 in CXCR4 due to the presence of a histidine at position 7.45. As a consequence of the pKa of about 4 for D2.50 in AT1h, the sodium pocket has a negative electrostatic potential (Fig. 2b) and should bind a sodium ion if this one can reach the allosteric site.

**Fig. 2:**
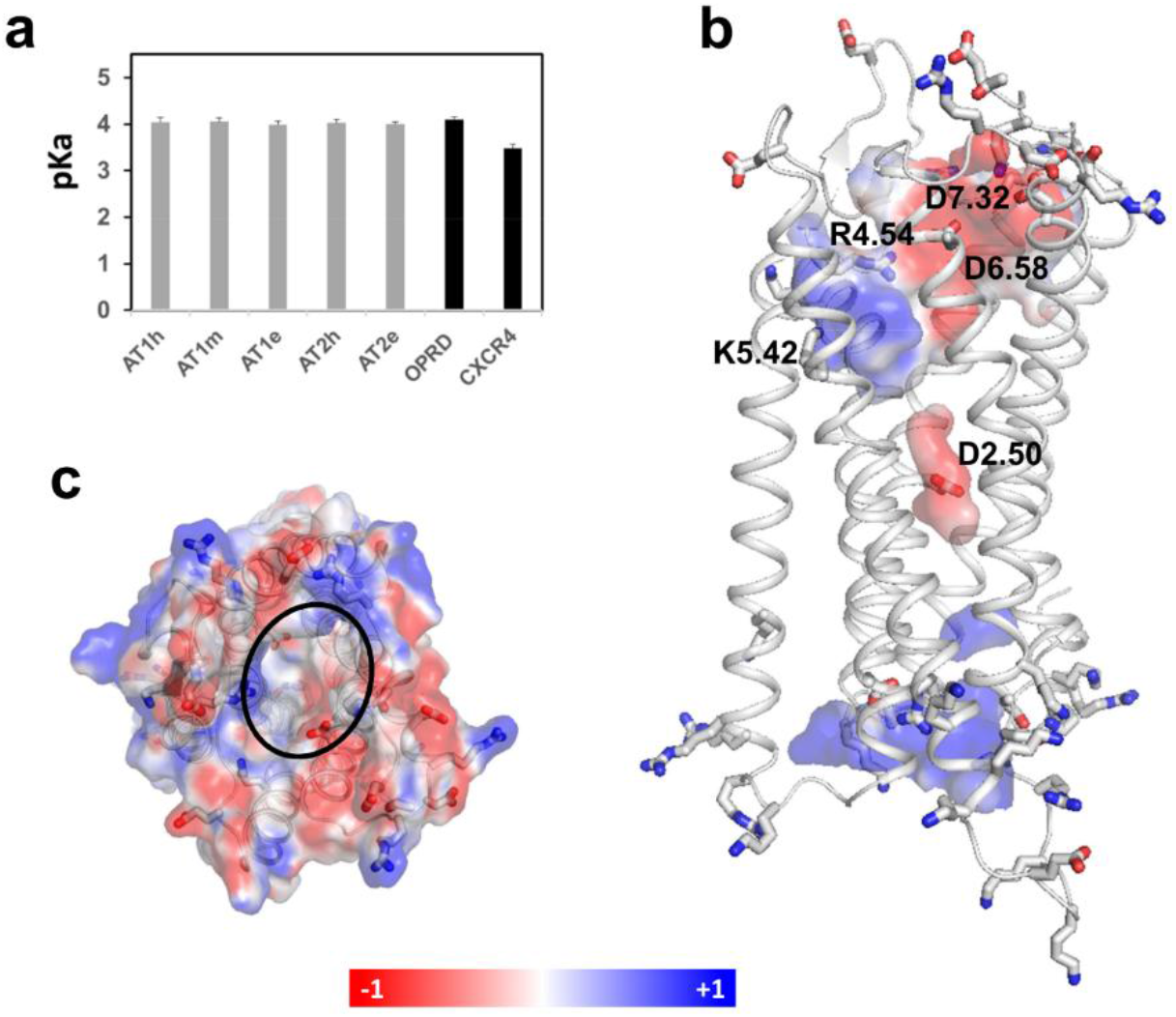
Electrostatic properties of AT1 and AT2. (a) pKas of residue D2.50 in AT1, AT2 and mutants. The pKas of D2.50 in CXCR4 and OPRD are given for comparative purpose; (b) Electrostatic potential of the extracellular surface of AT1h (top view); (c) Electrostatic potential of the AT1h internal cavities (side view). The charged side chains (Arg, Lys, Asp, Glu) are shown as sticks. The surfaces are colored from red (−1eV) to blue (+1eV). The electrostatic potential was computed with an ionic concentration of 0.15M NaCl.

### 3. Sodium ingress pathway

These findings prompted us to initiate a kinetics study to determine whether the sodium ion could reach the allosteric site. We modeled AT1h without sodium and embedded the receptor into a hydrated POPC bilayer in the presence of NaCl, both on the extracellular and intracellular water layers. These conditions allow testing the possibility of ingress from both the extracellular or the intracellular sides, since both pathways have been observed [31]. We used CXCR4 as control and we did observe fast sodium binding from the extracellular side (Supplementary Fig. S2) as observed by others [31]. Binding of a sodium ion to AT1h was not as easy and the same conditions (0.15M NaCl, cMD simulations) failed for AT1h in the submicrosecond timescale. Since our aim is the feasibility of sodium binding and not the details of the binding mechanism that can necessitate tens of microseconds [31], we decided to use harsher conditions by combining high sodium concentration (1M NaCl) and accelerated MD simulations in an attempt to speed up the kinetics.

After a 120 ns long cMD step, we initiated acceleration. In these conditions, we could observe a fast ingress of the sodium ion in the receptor internal cavity followed by its diffusion towards the allosteric pocket (Fig. 3 a,b). This fast step (<10ns) was followed by the desolvation of the ion from 5-6 to 2-3 water molecules in the first shell. Finally, after desolvation, the sodium ion was coordinated mainly to the D2.50 Oδ1 and Oδ2 atoms and to the N7.46Oδ1 atom (Fig. 3c), as observed under steady-state conditions (see below).

**Fig. 3:**
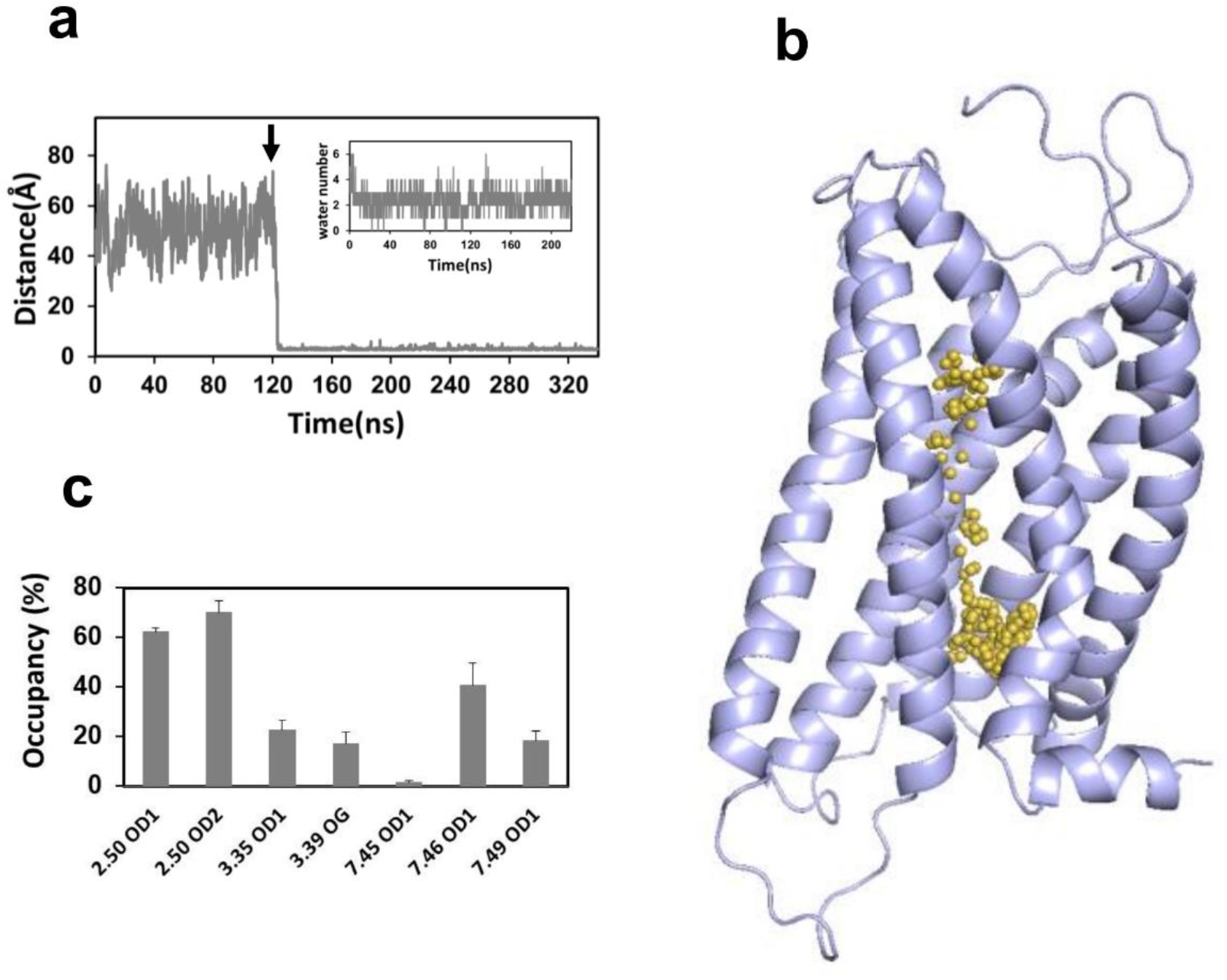
Kinetics of sodium binding to AT1h. (a) Trajectory of the sodium ion reaching the canonical binding site in AT1h, measured by the distance between the sodium ion and the Cγ atom of D2.50. The arrow indicates the beginning of the aMD simulations after 120 ns of cMD simulation. The insert represents the number of water molecules coordinating the sodium ion from the beginning of the acceleration; (b) Snapshots of the sodium trajectory (yellow spheres) during the first 20 ns of aMD simulations, superposed on the ribbon representation of AT1h (slate). The snapshots were taken every 0.16 ns; (c) Coordination of the sodium ion with protein atoms after desolvation (data are indicated as average ± standard deviations of three slots in the 20-220 ns range of the aMD trajectory).

The slow sodium ingress may be due to the complex electrostatic pattern of the extracellular surface of AT1 and to the high polarity of the orthosteric internal cavity (Fig. 2 b,c). This polarity is due to strictly conserved positive (K5.42 and R4.64) and negative (D6.58 and D7.32) residues that interact, respectively, with the terminal carboxyl group and the Arg2 sidechain of AngII in both AT1 [16] and AT2 [19]. A similar dipolar pattern is observed in AT1 and AT2 from both eel and human but not in CXCR4 (Fig. S3). In this latter receptor, the extracellular cavity possesses a negative potential which explains the fast sodium kinetics.

### 4. Sodium binding mode in the human angiotensin II receptors

We carried MD simulations of the human AT1 and AT2 receptors modeled in an inactive state, with a sodium ion located at the canonical site. In all AT1 and AT2 models, N3.35 was initially oriented inward, facing the sodium site, in the *trans* orientation, as observed in the inactive structures of AT1 [17]. In Fig. 4, we show typical examples of analysis for AT1h and AT2h trajectories, where we monitor sodium motion (RMSD), distances of the sodium ion to the putative coordination atoms and orientation of N3.35. All the analyses are reported in supplementary Fig. S4 and S5.

**Fig. 4:**
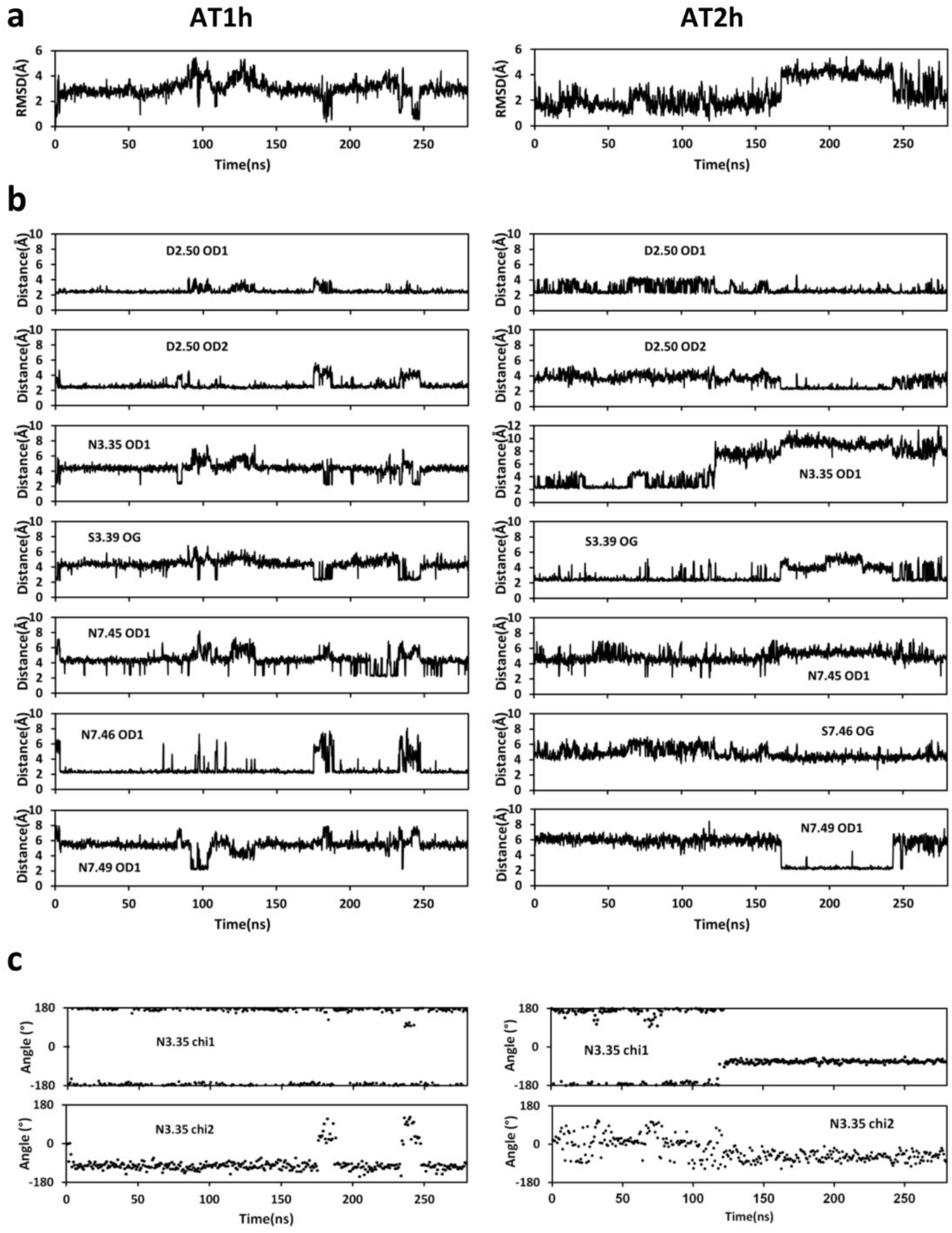
Stability of the sodium coordination in representative trajectories of AT1h (left) and AT2h (right). (a) RMSD of the sodium ion; (b) Distance of the sodium ion to putative coordination atoms; (c) Dihedral angles of N3.35 during the MD simulations.

In AT1h, the initial position of the sodium ion near N3.35 was not stable and the sodium rapidly moved by about 3-4 Å towards a position where it preferentially interacted with both the Oδ1 and Oδ2 atoms of D2.50, and the Oδ1 atom of N7.46, with only transient escapes towards N3.35, S3.39, N7.45 and N7.49. These interactions were similar to those observed for the sodium ion that reached the allosteric site from the extracellular compartment (Fig. 3). A typical snapshot of sodium coordination in AT1h is shown in Fig. 5a. The sodium ion is coordinated to the D2.50 Oδ1, D2.50 Oδ2 and N7.46 Oδ1 atoms and three water molecules. N3.35 does not interact directly with the sodium ion, but is located in the second shell. Its orientation is maintained by a conserved H-bond with D2.50 Oδ1 (Table II). By contrast, H-bonding with N7.46 is marginal.

**Table II.**
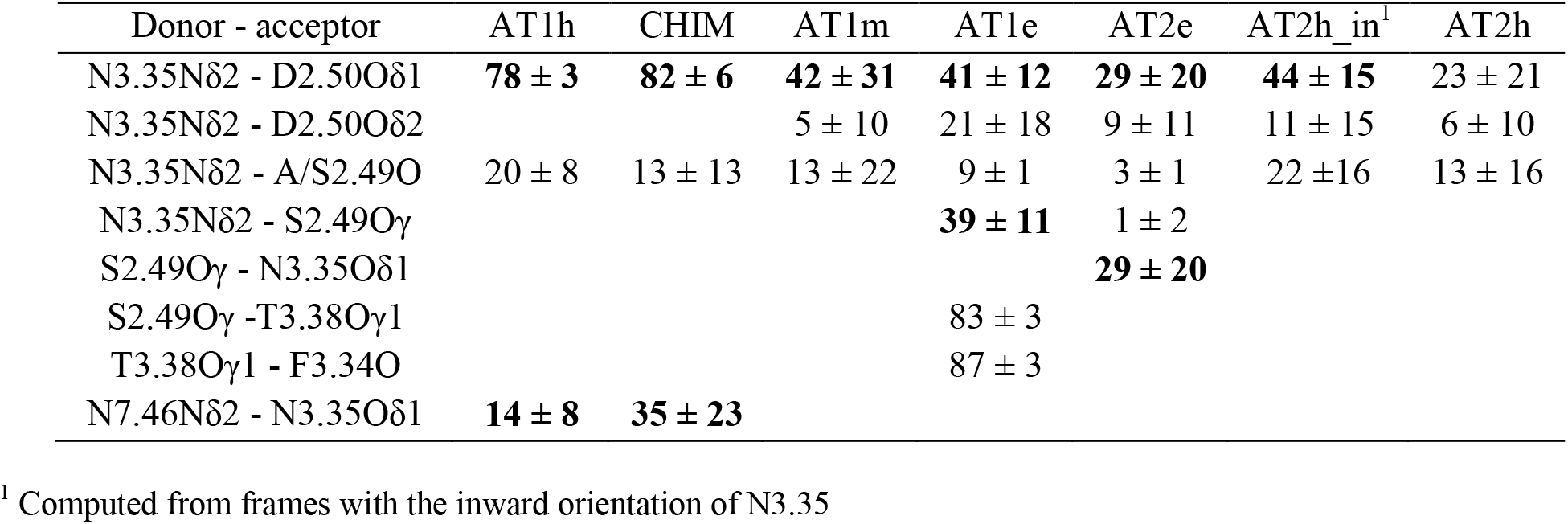
H-bonds stabilizing N3.35.

**Fig. 5:**
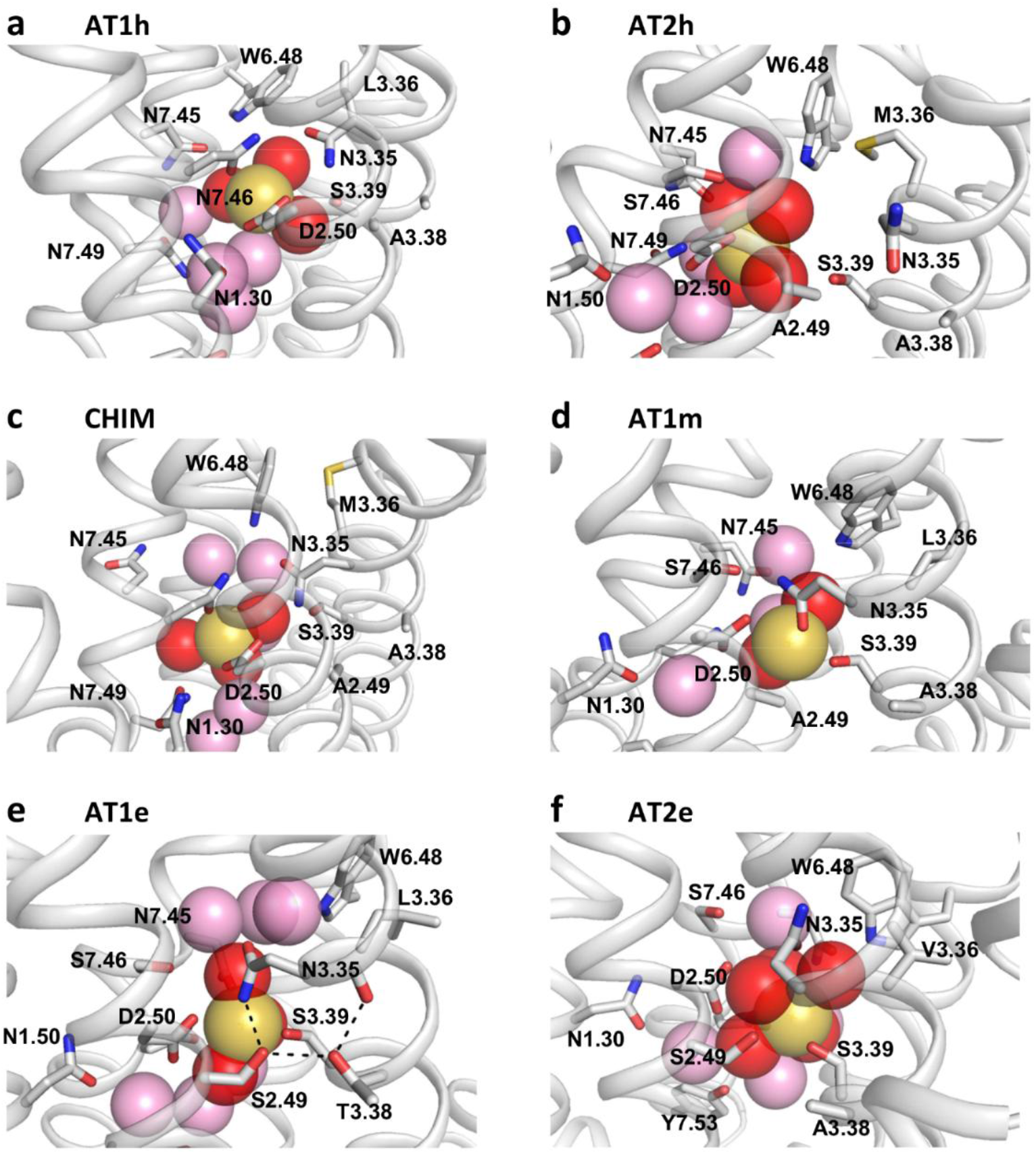
Snapshots of the sodium binding site in AT1 and AT2. (a) In the AT1h snapshot, the sodium ion is coordinated to D2.50 Oδ1 and Oδ2, to N7.46 Oδ1 and 3 water molecules; (b) In the AT2h snapshot, the sodium ion is coordinated to D2.50 Oδ1 and four water molecules. N3.35 is in the outward orientation; (c) In the CHIM snapshot, the sodium ion is coordinated to D2.50 Oδ2, N7.46 Oδ1 and 3 water molecules; (d) In the AT1m snapshot, the sodium ion is coordinated to D2.50 Oδ1, N3.35 Oδ1, S3.39 Oγ and two water molecules; (e) In the AT1e snapshot, the sodium ion is coordinated to D2.50 Oδ2, S3.39 Oγ and three water molecules. The dah lines indicate the H-bonds linking the N3.35 Nδ2, S3.39 Oγ, T3.38 Oγ and F3.31 O atoms; (f) In the AT2e snapshot, the sodium ion is coordinated to S3.39 Oγ and five water molecules. N3.35 is in the *g+* orientation (χ1 = 87°). D2.50 is in the second shell. In (a-f), the sodium ion is shown as a yellow sphere. Water molecules participating to the first coordination shell are shown as pink spheres. Other water molecules within 6 Å from the sodium ion are shown as pink spheres.

The sodium trajectory was markedly different in AT2h (Fig. 5). Initially, the sodium ion remained near the modeling position (1-2 Å) where it interacted with the D2.50 Oδ1, N3.35 Oδ1 and S3.39 Oγ atoms. However, sodium binding to N3.35 did not prevent outward rotamerization of N3.35. The N3.35 rotamerisation released the sodium ion which moved first towards N7.49 and then back toward S3.39 in an instable position where it was coordinated mainly to the D2.50Oδ1 atom and intermittently to the S3.39 Oγ atom. In the snapshot in Fig. 5b, the sodium ion is coordinated the D2.50Oδ1 atom and four water molecules.

The outward rotamerization of N3.35 was observed in 4 out of 6 trajectories carried out with two different AT2h model. The rotamerization occured at various times from 1 to 200 ns (Supplementary Fig. S2). These results support the assumption that the outward orientation of N3.35 is an intrinsic property of the human AT2 receptor.

### 5. Evolution of the sodium binding mode

The analysis of the sodium binding mode to human AT1 and AT2 was completed by the analysis of AT1 and AT2 from eels and of two mutants. The N7.46S AT1h mutant (AT1m) reverses the evolutionary observed S7.46N mutation whereas in the chimeric AT1h mutant (CHIM), a fragment of TM3 (3.25 to 3.40) was replaced by the equivalent sequence of AT2h. The CHIM mutant has a constitutive activity not observed for AT1 [32]. In all the initial models, the sodium ion was positioned near D2.50 and N3.35 as observed in the crystal structure of OPRD and N3.35 was oriented inward. A summary of the orientation of N3.35 during the simulations in the different replicas is reported in Table III.

**TABLE III.** Rotameric orientation of N3.35.

| Receptor | Number of runs | *trans* | *g-* | *g+* |
| --- | --- | --- | --- | --- |
| **AT1h** | 5 | **99.5 ± 1.0** | - | 0.5 ± 1.0 |
| **CHIM** | 3 | **99.8 ± 0.3** | - | 0.2 ± 0.3 |
| **AT1m** | 5 | **91.3 ± 19.3** | 8.6 ± 19.2 | 0.1 ± 0.2 |
| **AT1e** | 3 | **99.3 ± 0.9** | - | 0.7 ± 0.9 |
| **AT2e** | 3 | 75.2 ± 37.3 | - | **24.8 ± 37.3** |
| **AT2h** | 6 | 43.4 ± 43.3 | **56.1 ± 43.2** | - |

In most simulations, N3.35 remained oriented inward, in the *trans* conformation. In addition to AT2h, an outward rotamerisation was observed only in one out of five trajectories of the AT1m mutant after 150 ns. The *g+* conformation, where N3.35 points towards W6.48, was transiently observed in several trajectories, but was stable only in the AT2e receptor (Supplementary Fig. S5).

A summary of the sodium coordination in the various trajectories is shown in Fig. 6, while representative snapshots illustrating the variety of sodium coordination are reported in Fig. 5. Both in the AT1h receptor and in the CHIM mutant, the sodium ion bound preferentially D2.50 (two coordinations) and N7.46 (Fig. 6). In both cases, N3.35 remained in the *trans* orientation and strongly H-bonded with D2.50 (80% occupancy). Along with S3.39, it participated in the second coordination shell. A different binding mode was observed for the AT1m mutant. In this case, the sodium ion bound preferentially to D2.50 Oδ1, to N3.35 Oδ1 and to S3.39 Oγ atoms. This is similar to the binding mode observed in OPRD [33] and in the AT2h receptor with inward orientation of N3.35.

**Fig. 6:**
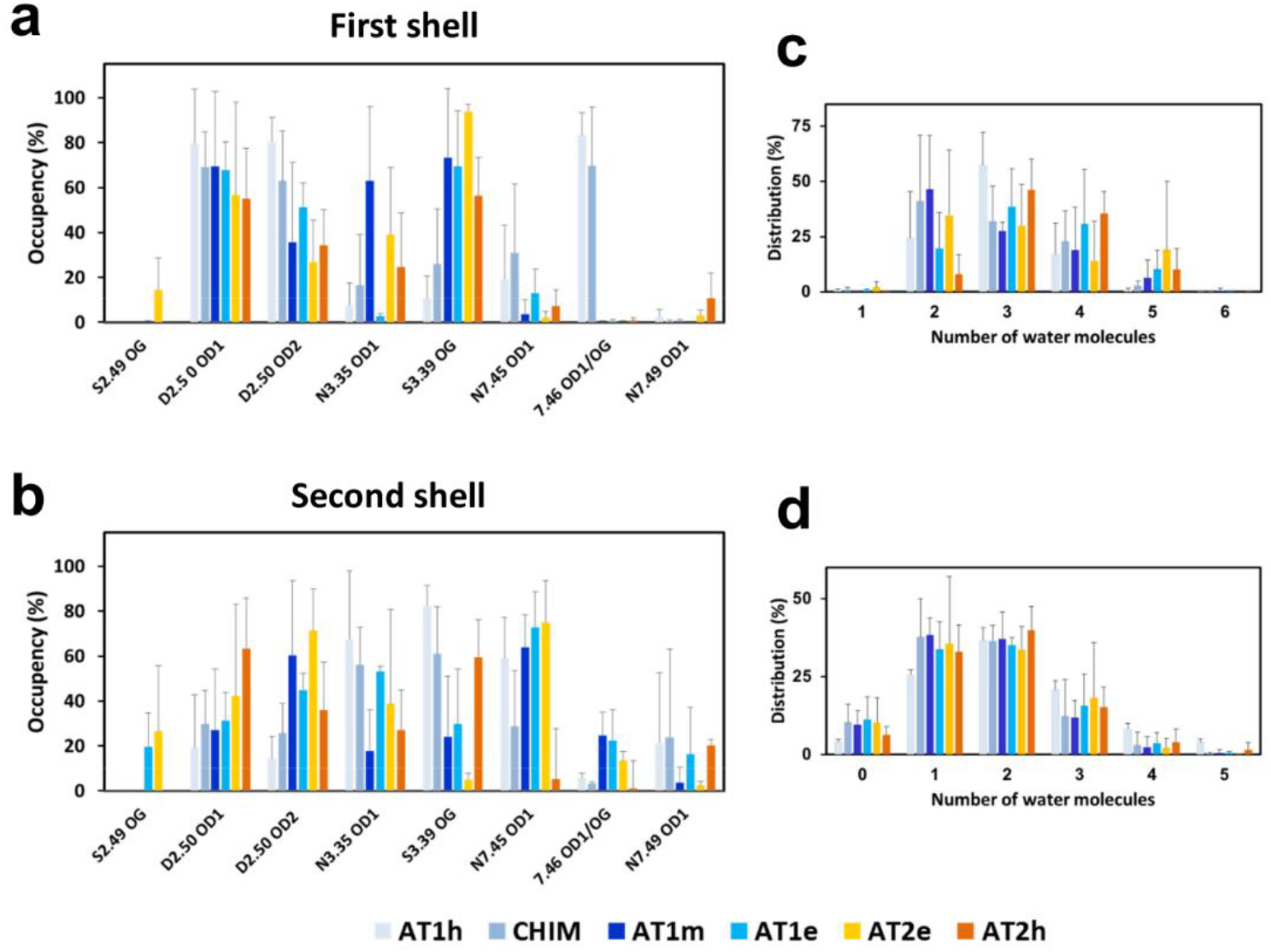
Coordination of the sodium ion in AT1 and AT2 receptors. (a, b) Average occupancy of the protein coordination atoms in the first shell (a) and in the second shell (b); (c, d) Distribution of water molecules in the first shell (c) and in the second shell (d). The receptors are indicated by the bar color code (AT1h: light blue, CHIM: middle blue, AT1m: dark blue, ATe: royal blue, AT2e: yellow, AT1h: orange).

**Fig. 7:**
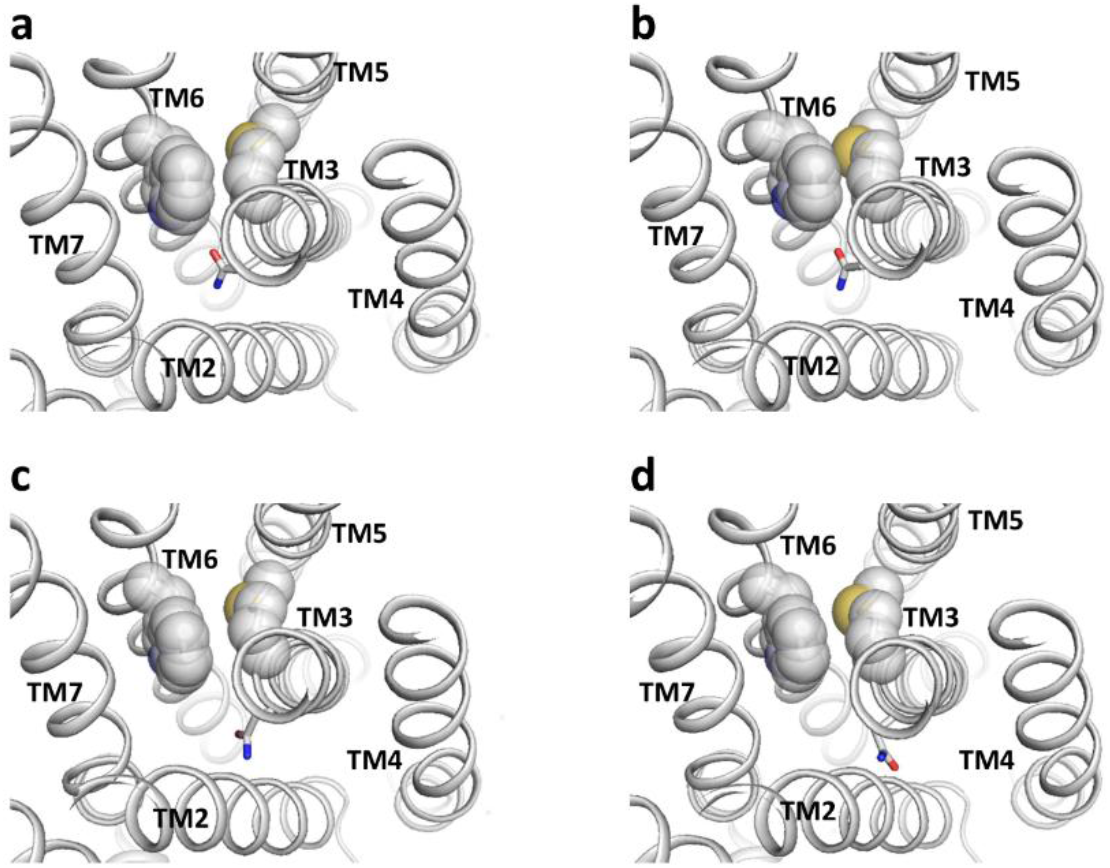
Rotamerization of N3.35 in human AT2. In (a), at t = 0, N3.35 is in the *trans* (χ_1_ = 161°) orientation. M3.36 is close to W6.48. In (b), at t = 8 ps, the M3.36 Sδ atom clashes with W6.48. In (c)m at t = 16 ps, M3.36 moves away from W6.48, the N3.35 sidechain initiates a rotational motion (χ_1_ = -146°). Finally, in (d), at t = 24 ps, N3.35 has rotamerized to the *g-*orientation (χ_1_ = -69°). In all the snapshots, the M3.36 sidechain is in the (*trans, g-*, t*rans*) orientation.

The AT1e receptor has the same S7.46 pattern as the N7.46S mutant but presents a different binding mode, with no coordination with N3.35 in the first shell. In this receptor, a network of polar interactions involving N3.35, D2.50, S2.49 and T3.38 (Table II) stabilizes the position of N3.35 at about 4 Å of the sodium ion (Fig. 5d). Thus, N3.35 does not participate to the first coordination shell but is part of the second shell (Fig. 6). Noteworthy, S2.49 and T3.38 are highly conserved (100% and 85%, respectively) in bony fishes, suggesting that this network is an important element of sodium allostery in bony fishs (Fig. 1).

In AT2e, S2.49 was not tethered to residue 3.38 (a strictly conserved Ala in AT2) and could participate in the coordination of the sodium ion along with Oδ1 or Oδ2 of D2.50, N3.35 and S3.39. This latter residue was always involved in the first shell whereas the other atoms could be involved either in the first or the second shell. In this receptor, N3.35 can be in the *g+* orientation (Table III) where it points towards W6.48 (Fig.5e). This orientation of N3.35 may be facilitated by the small size of the valine residue present at position 3.36 in AT2e. This position is usually a leucine in AT1 receptors and either a leucine or a methionine in AT2 receptors (Fig. 1).

The number of protein ligands in the first coordination shell varied from 2.8 ± 0.4 in AT1h to 1.9 ± 0.3 for AT2h. The first shell was completed by two to five water molecules. The second shell included around three protein ligands (varying from 2.5 ± 0.3 in AT1m to 3.1 ± 0.2 in AT1h and AT2h) and one to three water molecules (Fig. 6). In any case, four to five water molecules were within 6 Å of the sodium ion, with no significant differences between the receptors, indicating that the presence of an Asp residue at position 7.46 does not significantly reduce the size if the allosteric site.

## DISCUSSION

Most class A GPCRs possess a highly conserved sodium binding site. The best known effect of sodium is its role as negative allosteric modulator (NAM), with a strong stabilization of the inactive state. However, the role of the sodium ion is not limited to this “classical” NAM effect. Presence of sodium has dual effect on both reduced basal signaling and increased receptor stimulation by full agonists (reviewed in [22]). Some MD studies observed a transfer of a sodium ion through the membrane during receptor activation, due to the intracellular egress of the sodium upon activation [34]. Coupling of the sodium transfer from the extra to the intracellular side of the membrane is likely to make GPCRs sensitive to the electrostatic potential of the membrane [35,36]. Finally, mutations at the sodium binding site can induce biased or impaired signaling, suggesting a role as an allosteric cofactor of class A GPCR signaling [22].

In view of the putative sodium functions and of the role of residues within the sodium binding site of AT1 in the allosteric control of biased signaling [16,37], the question of sodium binding to the AT1 receptor is of primary importance to understand the activation mechanism of this receptor. To answer this question, we first deciphered the evolutionary history of the AT1 and AT2 receptors, with special focus on the sodium binding site (Fig. 1). While the sodium site in AT2 was maintained through evolution with few exceptions (Fig. 1), the sodium site of AT1 underwent an unusual S7.46N mutation during the transition to the terrestrian life suggesting that sodium binding might be impaired [16].

Arguments against maintained sodium binding rely on the absence of significant negative allosteric effects of sodium on receptor basal activity [38] and agonist affinity [16,39]. This might be due to H-bond interactions between N3.35 and N7.46 stabilizing the inactive state of AT1 in place of the sodium ion [16,37]. However, as detailed in a recent review [22], classical NAM effects on agonist binding have not been observed for at least one receptor (the β1 adrenergic receptor) whose sodium-bound structure has been resolved. Reversely, absence of sodium in crystallographic structures of putative sodium binding receptors is frequent.

Arguments supporting maintained sodium binding rely on several lines of evidence:

(1) The N7.46S AT1h mutant displays a constitutive activity at low but not at high sodium concentration. Such a behavior is usually interpreted as strong evidence for the stabilization of the inactive state by sodium binding [22]. This is consistent with the sodium binding mode of this receptor similar to OPRD (coordination to D2.50 Oδ1, N3.35 Oδ1 and S3.39 Oγ) [33].

(2) The pKas of the D2.50 residue in AT1 and AT2 receptors from humans and eels are similar to those computed for OPRD (around 4, see Fig. 2a). Consequently, for all the receptors investigated, D2.50 should be negatively charged at neutral pH. The sodium cavity is large enough to contain at least four or five water molecules (Fig. 5 and 6), indicating absence of steric hindrance to sodium binding in AT1h.

(3) A sodium ion can actually reach its binding site in human AT1 from the extracellular site (Fig. 3). The dipolar electrostatic potential of the orthosteric cavity is due to highly conserved charged residues in AT1 and AT2 and is similar in all receptors investigated (Supplementary Fig. S3), supporting similar sodium ingress mechanism.

Taken together, these results strongly support the assumption that a functional sodium binding site has been maintained in both AT1 and AT2 receptors throughout evolution. However, the sodium binding modes markedly differ between eels and humans for both AT1 and AT2, in link with hallmark mutations in the evolution of these receptors during the transition to terrestrian life.

Fish receptors possess a serine at position 2.49. The presence of either Ala or Ser at this position is frequent in GPCRs. However, the presence of a threonine at position 3.38 facing a serine at position 2.49 as observed in fish AT1 is unusual (only three occurrences in human GPCRs). This pattern stabilizes the sodium binding site through an H-bond network linking N3.35, S2.49 and T3.38 (Table II). Its high conservation in fish AT1 (Fig. 1b) suggests that it is an important element of sodium allostery and tight control of receptor activation. This pattern is not observed in fish AT2 but N3.35 is still engaged, to a lesser extent, in H-bonds with S3.39..

In amniota AT1, the destabilization due to the S2.49A mutation is compensated by the S7.46N mutation. N3.35, in the second coordination shell, is stably H-bonded to D2.50 (around 80%) and maintained in the inward orientation. This stable H-bonding pattern is also observed in the CHIM mutant, but not for the N7.46S mutant, which may facilitate the outward rotamerization of N3.35 (Table III).

An important consequence of the S7.46N mutation in amniota AT1 can be inferred from the promiscuity of the human N7.46S AT1 mutant [38]. In particular, the N7.46S mutant can be activated by Ang1-7, AngIV, and Ang5-8. Quoting Feng, “The promiscuous agonist specificity displayed by N295S […] in this study could be disastrous if a similar mutation were to occur in AT1 receptors in nature” [38]. Actually, in nature, this is the reverse mutation which occurred in amniota (Fig. 1b), preventing AT1 activation by these peptides. In addition, the versatility of the interactions in which N7.46 can be involved makes AT1 prone to biased signaling. Indeed, the resolution of AT1 in complex with AngII and β-arrestin-biased ligands, along with extensive MD simulations, have shown that different active conformations can be stabilized by different orientations and H-bonding patterns of N3.35 and N7.46 [16,37].

Concerning human AT2, full characterization of this receptor is difficult because of absence of response in conventional GPCR essays [4,13]. However, the pharmacological profiles based on binding affinities reveal that AT2 lacks the specificity of AT1 for AngII as compared to either synthetic or natural AngII derivatives [21,40,41]. The promiscuous binding of AT2 to AngII derivatives suggests a “relaxed” conformation by analogy with constitutive AT1 mutants [21]. This assumption is supported by the crystal structures of AT2 in active forms only [18-20]. In these structures, N3.35 is in the outward orientation. The rotamerization of N3.35 from the inward to the outward orientation was observed in several AT2h trajectories (Table III). The outward orientation favors sodium release and subsequent activation [42]. Noteworthy, in one of these trajectories, we observed a steric clash between W6.48 and M3.36, just before the N3.35 outward motion, suggesting that the presence of a methionine at position 3.36 might favor the rotamerization of N3.35.

An outward N3.35 rotamerization was not observed in eel AT2 (Table III), supporting the assumption of a tighter sodium binding site in this receptor as compared to human AT2. Two elements might contribute to it: (1) the presence of a serine rather than an alanine at position 2.49 which increases the H-bonding pattern (Table II) and (2) the presence of a valine rather than a methionine at position 3.36. This position is involved in receptor activation upon AngII binding both in human AT2 [19] and in human AT1 [16]. The bulkier M3.36 in human AT2 as compared to L3.36 in human AT1 may facilitate activation of AT2 by AngII derivatives missing the bulky terminal Phe-8 [19]. Among them, Ang1-7, which can bind to and activate human AT2 [41,43], does not activate eel AT2 [44].

The transition from aqueous to terrestrial environment was accompanied by major changes in the renin-angiotensin system. The conventional activation of a G protein by AT2, observed in eels [44], has been lost in mammalians [4,13] whereas novel receptors, such as MAS, emerged during the transition [45] and participate in the counter-regulatory arm of the RAS [1]. MAS receptor is activated by Ang1-7. We can note that Ang1-7 became an agonist of AT2 [41,43] and a β-arrestin biased agonist of AT1 [46,47] in amniota, after MAS emergence. Taken together, the key mutations observed in amniota AT1 and AT2 (Fig. 1) participate in the split between the classical (ACE/AngII/AT1) and counter-regulatory (ACE2/Ang1-7/MAS, AT2) RAS pathways [48].

It is worth to note that the development of the counter-regulatory axis of the RAS is correlated with the major evolution of the cardiovascular system during the transition to terrestrial life [49]. Fishes have a single circulatory system with a two chamber heart. Terrestrian life requires uptake of oxygen from air, which led to the emergence of lungs, of two circulatory systems and of a heart with three (amphibian) or four chambers (amniota). The two circulatory systems have very different blood pressure, in particular for amniota which need high pressure in systemic circulation to fight gravity and low pressure in lungs to favor oxygen exchange [49]. The key mutations in AT1 and AT2 summarized in Fig. 8 arose at that moment and might have helped an artful balance between the two arms of the renin-angiotensin system.

**Fig. 8:**
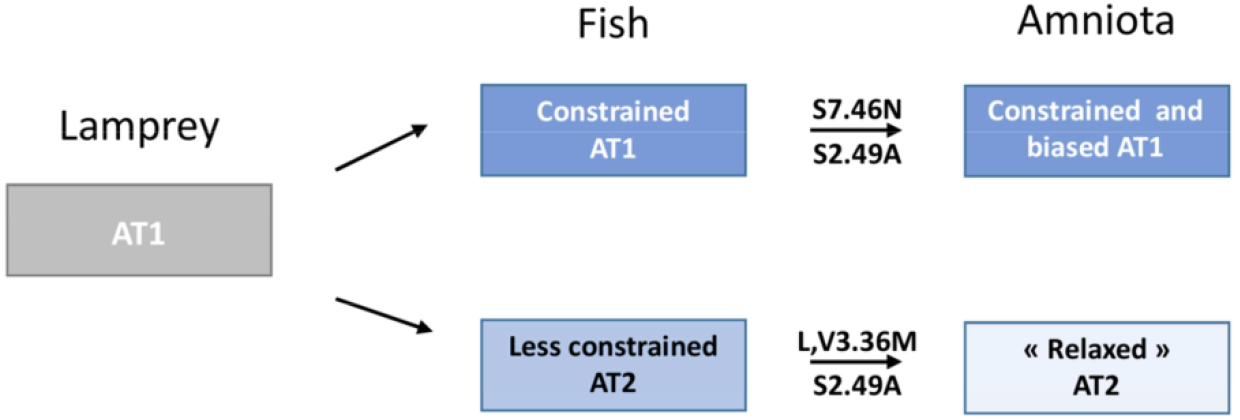
Schematic representation of the evolution of the Angiotensin II receptors.

## MATERIALS AND METHODS

### Sequence retrieval, clustering and analysis

Angiotensin receptor sequences were obtained from Uniprot (www.uniprot.org) [50]. The initial exhaustive request using the keywords “IPR000276” (Interpro identifier for class A GPCRs) and “angiotensin” led to 550 protein sequences. To eliminate false positives or truncated sequences, a multiple sequence alignment was carried out with ClustalX [51], and then a manual selection was carried out using GenDoc [52] to obtain a set of 318 “clean” sequences.

The over representation of highly similar mammalian sequences in this set (120 mammalian AT1 sequences out of 203, 68 mammalian AT2 sequences out of 120) prompted us to build a non-redundant set using the perl script nrdb90.pl [26]. This program builds clusters with a sequence identity threshold of 90%. This approach led to 38 AT1 and 37 AT2 clusters. Among them, there were 3 and 5 mammalian clusters for AT1 and AT2, respectively. The clusters of human AT1 and AT2 included 115 and 39 sequences, respectively. A representative sequence was selected for each cluster, yielding a non-redundant set of AT1 and AT2 receptors. It is noteworthy that there is no sequence of AT2 from cartilaginous fishes reported in Uniprot. To determine whether the absence of AT2 sequences was due to a sequencing problem or to the evolutionary difference, we extended the search to AT2 sequences of the whale genomes in Genbank. In the two additional genomes reported, we found AT2-like sequences but they should be nonfunctional, either because of an AngII binding impairing K5.42I mutation (*Rhincodon typus*) or an N-terminal truncation (*Chiloscyllium punctatum*). No other example of such mutations was observed in the entire set of reported AT2 sequences, supporting the assumption that functionnal AT2 gene was lost in cartilagineous fishes.

The representative sequences of the clusters were aligned with ClustalX and then a Neighbour-Joining (NJ) phylogentic tree of the representative sequences of the clusters was built with the MEGA5 software [53] using 500 replicates for bootstrapping. Logo plots were obtained from the Weblogo site [54].

### Molecular modeling

The inactive state of human AT1 (AT1h) with bound sodium was modeled from the sodium free 4YAY [17] and 4ZUD [39] structures of AT1 and from the sodium bound 4N6H structure of the δ opioid receptor (OPRD) structure [33], using the homology modeling software MODELER [55]. The sodium bound AT1h was modelled from residue 17 to 317 with 7 water molecules in the sodium pocket positioned by homology with 4N6H. The C-terminal helix 8 (H8) has been modeled by homology with 4NH6 because in the 4YAY and 4ZUD structures, helix 8 (H8) is either missing or bent relative to the membrane plane. The bent orientation of H8 leads to swing-saw motion of this helix and transmembrane helix 7 during MD simulations [56]. The apo model of AT1h without bound sodium was obtained by replacing the sodium ion by a water molecule. The model of the N7.46S AT1h mutant was prepared from the model of AT1h using the Charmm Gui interface [57]. The model of the chimeric CHIM mutant was prepared using MODELLER from the model of AT1h by substituting the sequence from residue 3.25 to 3.40 by the sequence from equivalent positions in AT2h [32]

~~~
AT1h: CKIASASVSFNLYASV
AT2h: CKVFGSFLTLNMFASI
~~~

Two models of inactive human AT2 were built using MODELLER from the active-like AT2 structure 5UNG [20] and the inactive AT1h model. The intracellular halves of TM5, TM6 and TM7 and the C-terminal H8 from the 5UNG structure were truncated and replaced by the equivalent parts of the AT1h model, including the sodium ion and the water molecules. The two models differ in the length of TM6. In the first model (AT2ha), TM6 had the same length as in AT1h. In the second model (AT2hb), α-helical restraints were added to the modeling procedure to insure that TM6 length was identical to the length in the 5UNG structure. In either case, the AT2h models were built from residue 35 to 333 and contained a sodium ion and 7 water molecules in the sodium pocket. In addition, N3.35 was modeled in the *trans* orientation as observed in the inactive AT1 structures (PDB 4YAY and 4ZUD) and faced the sodium binding site.

Models of sodium bound AT1 and AT2 from Japanese eel (AT1e and AT2e, respectively) were built by homology with the models of AT1h and AT2hb.

### Molecular dynamics simulations

The models were prepared for molecular dynamics simulations (MD) using the Charmm-gui interface [57]. The models were embedded in a palmitoyl-oleoyl-phosphatidyl-choline (POPC) bilayer with 60 lipids on each layer and solvated in a TIP3P model for water molecules with all atoms represented explicitly. For simulations aimed at analyzing the sodium binding mode in models with a sodium ion in the allosteric site, the charges were neutralized by adding chloride ions. For simulations aimed at analyzing the sodium pathway towards the binding pocket, the sodium ion was replaced by a water molecule in the AT1h model. The charges were neutralized by adding either 0.15 M or 1M NaCl. In all the simulations, the D2.50 residue located within the sodium binding site was negatively charged, in agreement with the pKa of this residue (see below) that was calculated with DelphiPKa [58].

Molecular dynamics simulations of the receptor models embedded into the hydrated POPC bilayer were carried out using NAMD v2.9 MD software [59] and the CHARMM36 parameter set [60,61]. They were performed using the HPC resources of IDRIS, granted by GENCI (www.genci.fr). The entire assembly was subjected to energy minimization for 5000 steps to remove close contacts between receptor atoms and solvent or lipid layers. Equilibration of protein–membrane system was carried out with a modified version of a protocol developed elsewhere [25]. The equilibration protocol included first six steps in which harmonic restraints were gradually taken off to achieve a smooth relaxation, for a total of 1 ns, and then a 20 ns step carried out under the same conditions as the production run to achieve stable conditions. In the first two equilibration steps, the NVT ensemble at 310 K and time step of 1 fs were used. The following equilibration and production steps were carried out at constant temperature (310 K) and pressure (1 atmosphere), using a 2 fs time-step for integration. The Particle Mesh Ewald method (PME) was used to calculate the electrostatic contribution to non-bonded interactions with a cutoff of 12 Å. The cutoff distance of the van der Waals interaction was 12.0 Å. The SHAKE algorithm was applied to the system. Each trajectory lasted 280 ns (20 ns for equilibration and 260 ns for production).

The dual boost accelerated molecular dynamics simulations were carried out as implemented in NAMD [62]. This technique, based on an extended biased potential MD approach, efficiently enhances conformational sampling. The dual boost approach works by adding (1) a dihedral potential boost to all dihedral angles in the system, which affects protein atoms, and (2) a total potential boost to all atoms in the system, which affects also water and ions, when the energies are below a threshold [63]. In the soft protocol that we use [25], the threshold energies are set to the average energies, *E*_*dihed_avg*_ and *E*_*pot_avg*_, computed from classical MD and the acceleration factors are calculated according to:

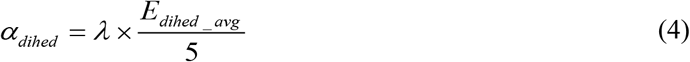

and

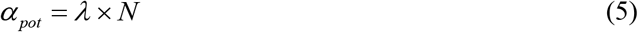

where the acceleration parameter λ is set at 0.3 and *N* represents the number of atoms in the system. The 20-120 ns range of classical MD simulations was used to calculate the acceleration parameters. The snapshot obtained after 120 ns was used to initiate the accelerated simulation

Trajectories were graphically analyzed with VMD [64] and PYMOL (DeLano Scientific LLC, San Franscisco, USA). Quantitative analyses were carried out with home-developed scripts written in tcl/Tk for the VMD software and in R using the Bio3D R package [65]. H-bonds were measured with the H-bond facility of VMD with cutoffs of 30° and 3.5Å. PYMOL was used for figure preparation.

### Electrostatics potential and pKa computations

Electrostatics potentials were computed by a Poisson-Bolzmann approach with the Delphi program using the smooth Gaussian dielectric function developed by Alexov and al [28]. Similarly, the pKa values of the receptor ionizable residues were computed with DelphiPKa using the same Gaussian dielectric function [29,30]. Both Delphi and DelphiPKa have been installed locally. In the Gaussian representation of atomic density resulting into a smooth dielectric function, two parameters have to be defined: the width of the Gaussian function, σ, and the internal permittivity of the protein, ε_in_. The default values for external ionizable groups are ε_in_ = 8 and σ = 0.7, whereas the default values for internal groups are ε_in_ = 4 and σ = 0.9. In these studies, we aimed at determining the electrostatic properties of D2.50, located at the sodium binding pocket, in AT1 and AT2. The sodium pocket is lined by several polar groups, including the aspartic acid D2.50, and is wide enough to contain a sodium ion and several water molecules. To determine adequate parameters, we compared the pKa of D2.50 in humain AT1 and in the δ opioid receptor (OPRD), a GPCR with resolved crystal structure in a sodium bound inactive state [33], as a function of a variety of ε_in_ and σ values. For each pair, analyses were done on seven equally spaced frames from MD trajectories. All the combinations of ε_in_ and σ values led to similar pKa values for D2.50 in OPRD and AT1h. (Supplementary Fig. S1). Most of them led to a pKa value < 6, consistent with deprotonation of D2.50 in physiological conditions for both receptors. This is not the case for the default values for internal groups that led to a value of 7.3 ± 1.0 for the pKa of D2.50 in OPRD, which is not consistent with the observed affinity of the sodium ion for OPRD [22]. By contrast, the default values for external ionizable groups led to a D2.50 pKa value of 4.1, which supports a negatively charged aspartic acid interacting with the sodium ion. Consequently, the latter set (ε_in_ = 8 and σ = 0.7) was used both for electrostatic potentials and pKa calculations. For each receptor investigated, pKa computations were done on a representative set of seven frames from MD trajectories.

The electrostatics potentials were calculated with the Gaussian-based method on the receptors placed at the center of a cube filled with an aqueous medium, implicitly modeled by a dielectric permittivity ε_ex_ of 80. The axes of the cube are oriented along the principal axis of the solute. The edge size of the cube, L, depends on the maximal dimension of the solute Lp (Lp = L * 0.7). The space was discretized with 2 grid points by Å. The permittivity of the receptor ε_in_ was 8 and the Gaussian width, σ, was 0.7. The ionic strength was set at 0.15 M. The atom charges and sizes used for the calculations were based on the default Charmm parameters provided by Delphi (Charmm22).

## Acknowledgments

We thank Dr Guillaume Lebon (Institut de génomique fonctionnelle, Montpellier, France) for critical reading of the manuscript and stimulating discussion. This study was supported by institutional grants from INSERM, CNRS and University of Angers. This work was granted access to HPC resources of IDRIS (GENCI grant 100567 to MC). MC is supported by CNRS. AT is supported by a fellowship from the University of Carthage (Tunisia). RB is supported by a fellowship from the University of Angers (France).

## Supplementary Data

Fig. S1: Comparison of D2.50 pKa in OPRD and AT1 as a function of delphi parameters.

Fig. S2: Comparison of the binding kinetics of the sodium ion to CXCR4 and AT1.

Fig. S3: Comparison of the electrostatics potentials of the internal cavities of human and eel AT1 and AT2 receptors, the CHIM and N7.46S AT1 mutants, and of CXCR4.

Fig. S4: Distances of the sodium ion to putative protein ligands in the MD simulations of human and eel AT1 and AT2 receptors, and the CHIM and N7.46S AT1 mutants.

Fig. S5: Rotamers of N3.35 in the MD simulations of human and eel AT1 and AT2 receptors, and the CHIM and N7.46S AT1 mutants.

