## Supplementary Data for "Molecular evolution of the Angiotensin II receptors AT1 and AT2: Specificity of the sodium binding site in amniota"

Fig_S1: Comparison of D2.50 pKa in OPRD and AT1 as a function of delphi parameters.

Fig_S2: Comparison of the binding kinetics of the sodium ion to CXCR4 and AT1.

Fig_S3: Comparison of the electrostatics potentials of the internal cavities of AT1 and AT2 from eel and human, the CHIM and N7.46S AT1 mutants and of CXCR4.

Fig_S4: Distances of the sodium ion to putative protein ligands in the MD simulations of AT1 and AT2 from human and eel, and the CHIM and N7.46S AT1 mutants.

Fig. S5: Rotamers of N3.35 in the MD simulations of AT1 and AT2 from human and eel, and the CHIM and N7.46S AT1 mutants.

### Fig. S1

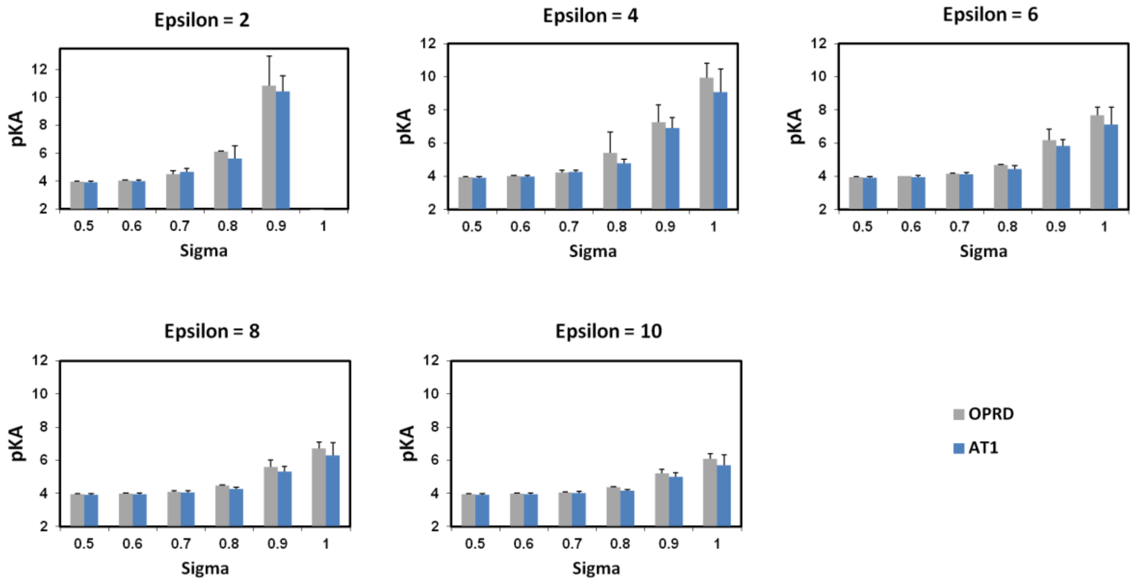


**Fig. S1: Comparative analysis of the pKa of D2.50 in OPRD and human AT1 as a function of Delphi parameters.** pKa were computed with DelphiPKa using the Gaussian representation of atomic density that depends on the internal Epsilon and on the width of the Gaussian function, Sigma. For each pair of parameters, analyses were done on seven equally spaced frames from MD trajectories.

### Fig. S2

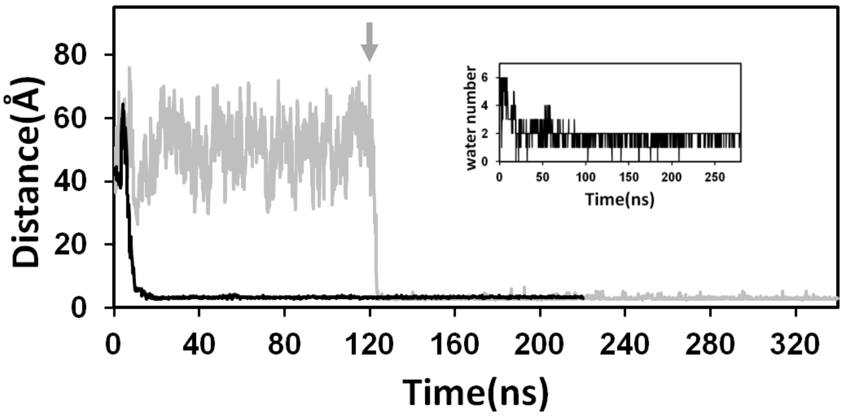


**Fig. S2:** Comparison of the binding kinetics of the sodium ion to CXCR4 (black) and AT1 (grey). The grey arrow indicates the acceleration of the AT1 trajectory. The insert represents the desolvation of the sodium ion in CXCR4. Classical MD simulations of CXCR4 have been carried out with sodium free CXCR4 embedded into a hydrated POPC membrane. The sodium chloride concentration was 0.15M.

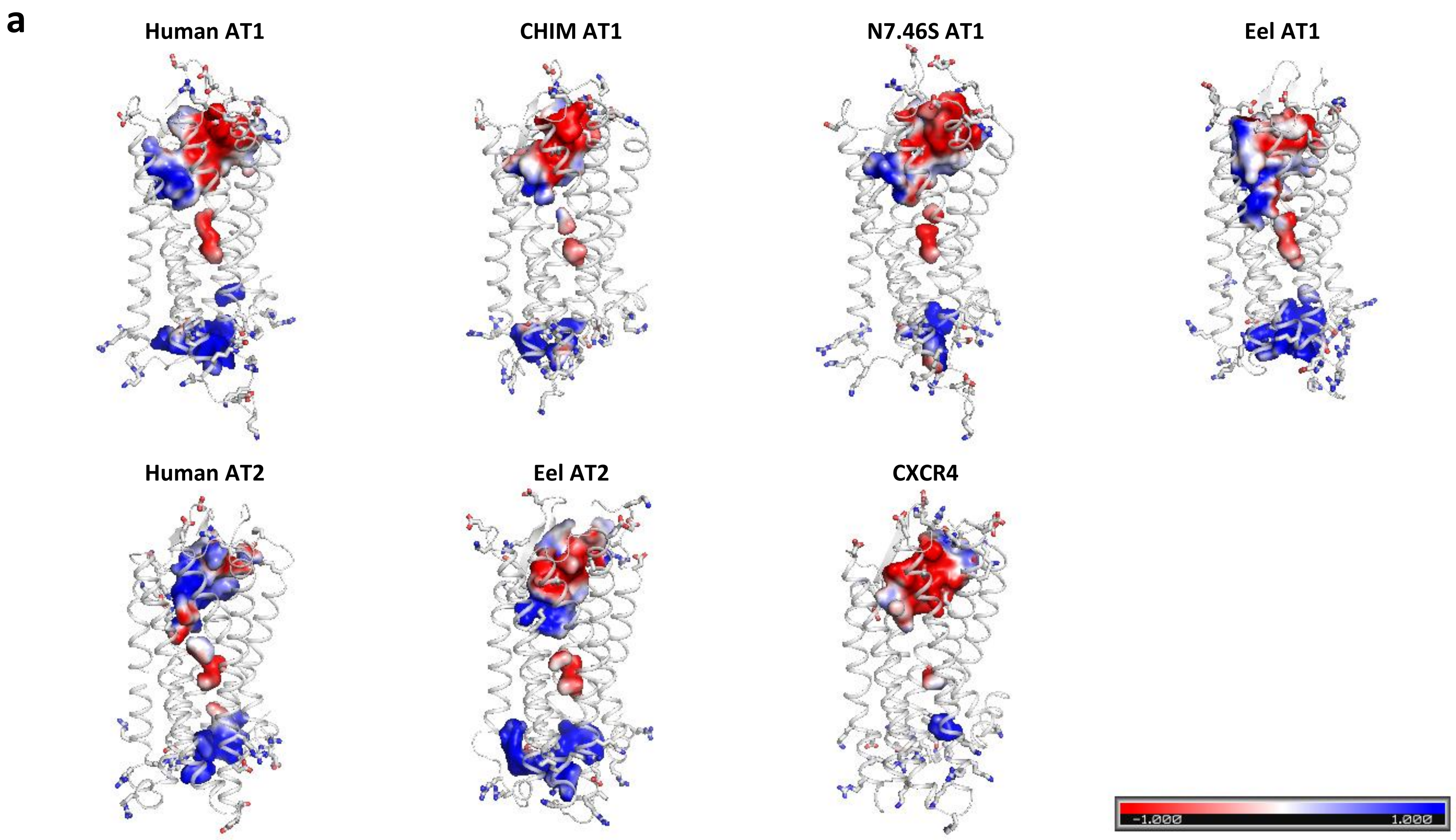
a
Human AT1
CHIM AT1
N7.46S AT1
Eel AT1
Human AT2
Eel AT2
CXCR4
-1.000
1.000

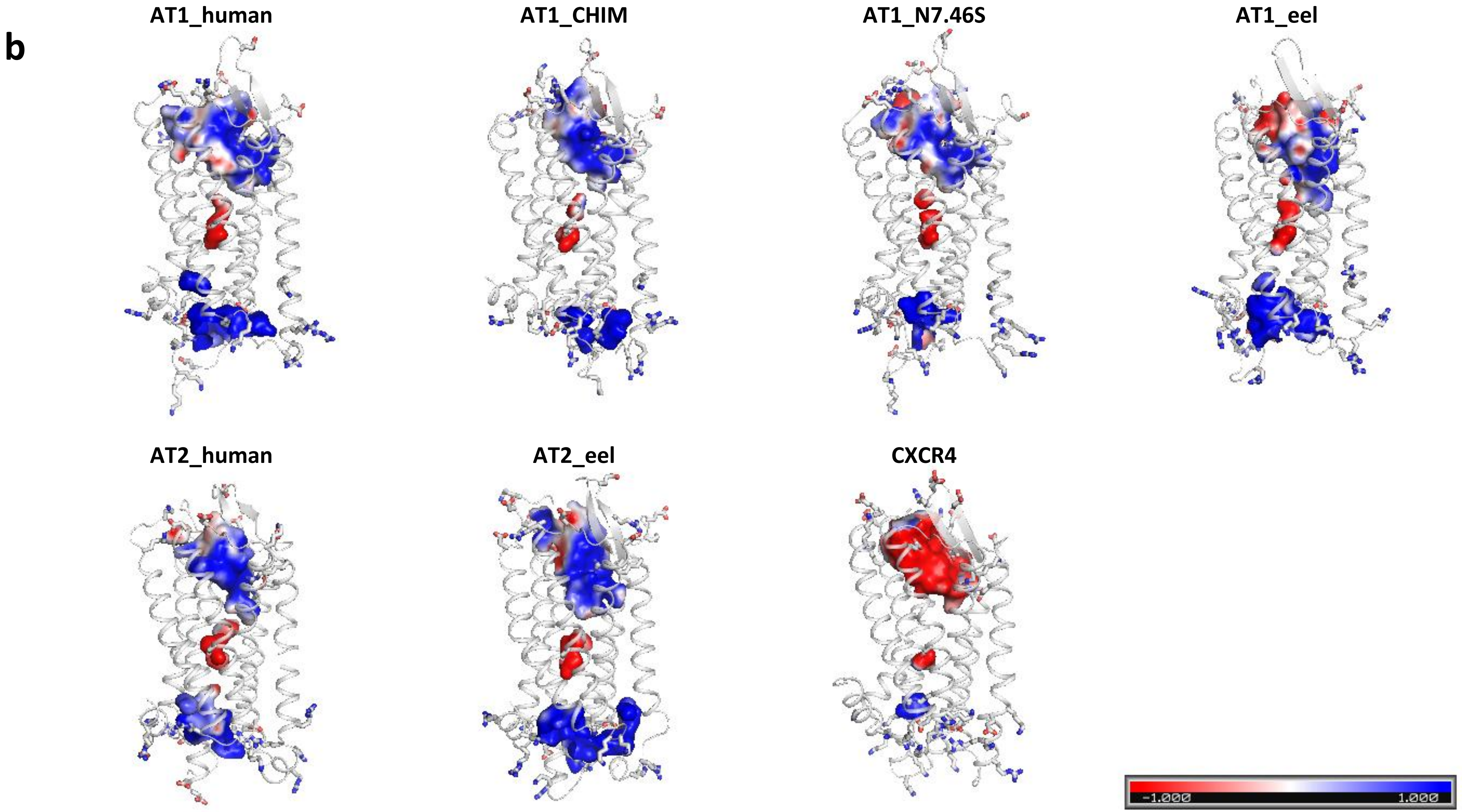


**Fig. S3:** Electrostatic potential of the receptor internal cavities in human and eel AT1, human and eel AT2, the CHIM and N7.46S AT1 mutants and CXCR4, given for comparative purpose. (a) and (b) are two views from the lateral side, differering by a 180° rotation.

**Fig_S4:** Distances of the sodium ion to putative protein ligands in the MD simulations of AT1 and AT2 from human and eel, and the CHIM and N7.46S AT1 mutants.

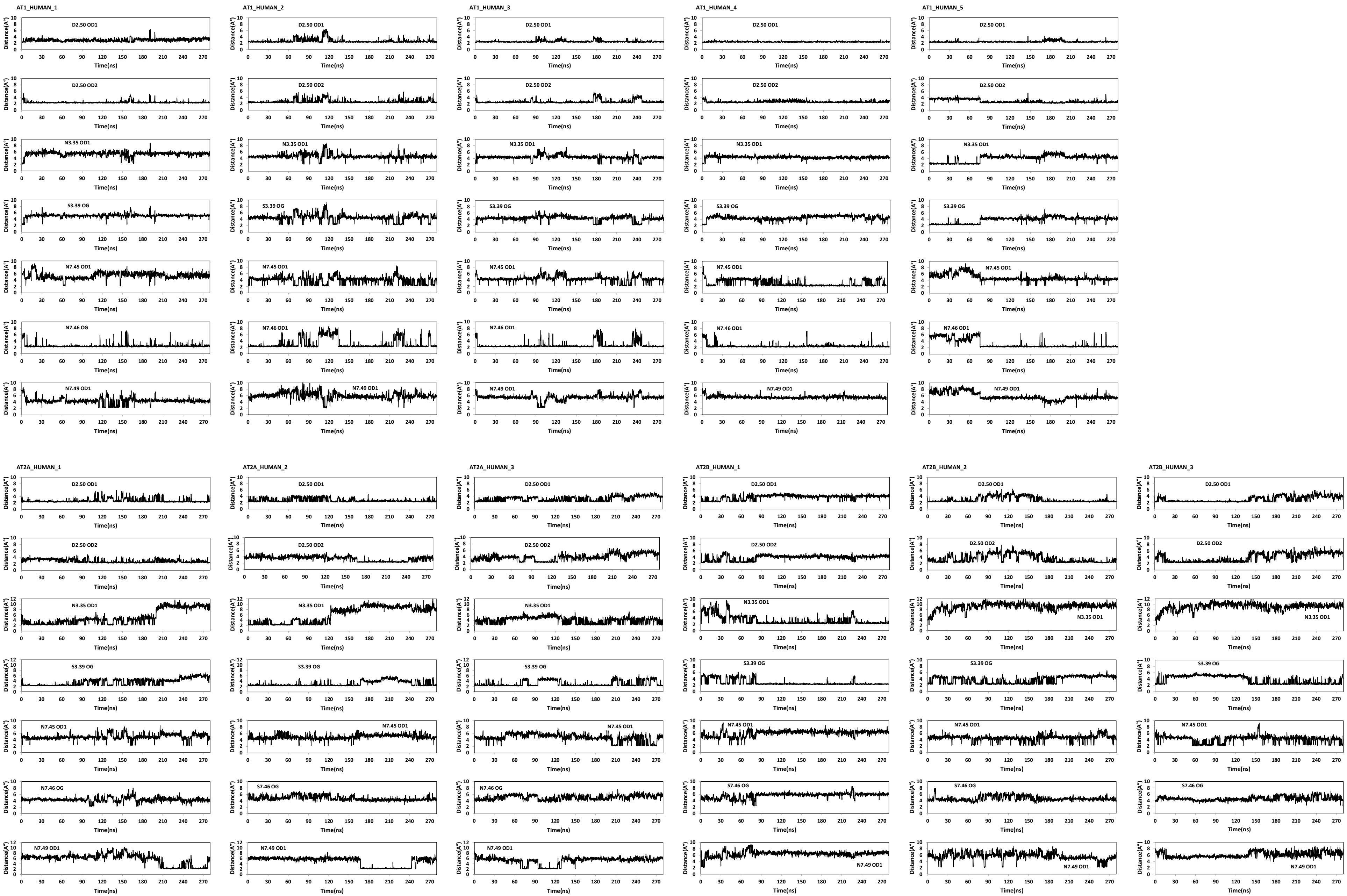
AT1_HUMAN_1
AT1_HUMAN_2
AT1_HUMAN_3
AT1_HUMAN_4
AT1_HUMAN_5
D2.50 OD1
D2.50 OD2
N3.35 OD1
S3.39 OG
N7.45 OD1
N7.46 OG
N7.46 OD1
N7.49 OD1
Distance(A°)
Time(ns)
AT2A_HUMAN_1
AT2A_HUMAN_2
AT2A_HUMAN_3
AT2B_HUMAN_1
AT2B_HUMAN_2
AT2B_HUMAN_3
S7.46 OG

AT1_N7.46S_1

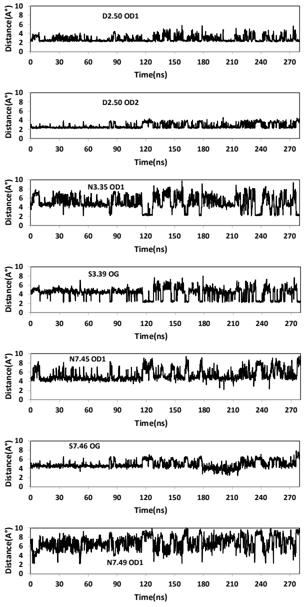


AT1_N7.46S_2

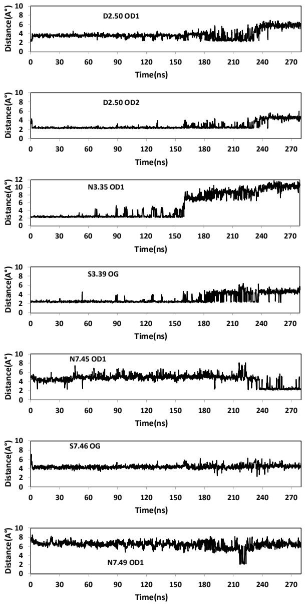


AT1_N7.46S_3

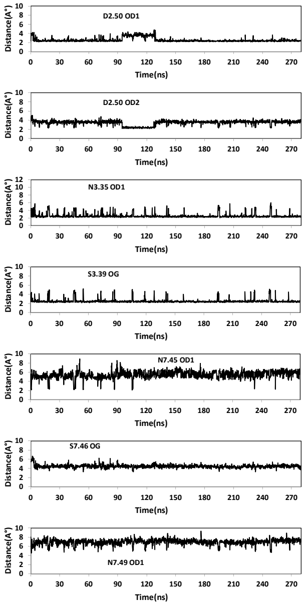


AT1_N7.46S_4

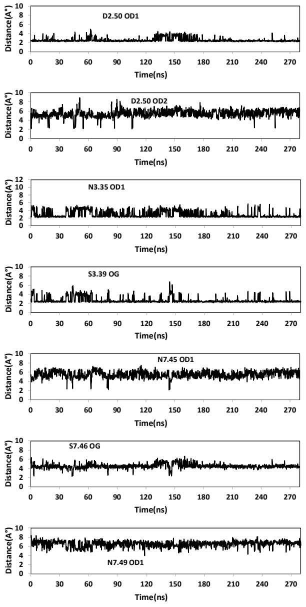


AT1_N7.46S_5

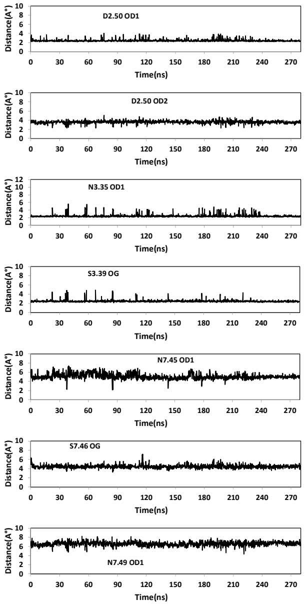


AT1_CHIM_1

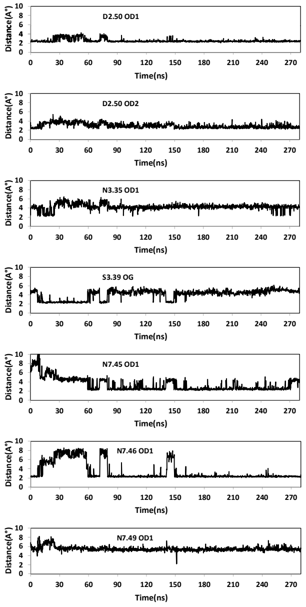


AT1_CHIM_2

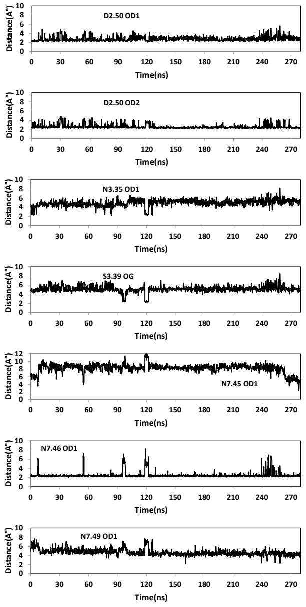


AT1_CHIM_3

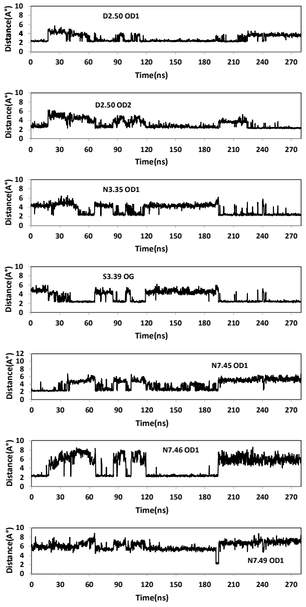

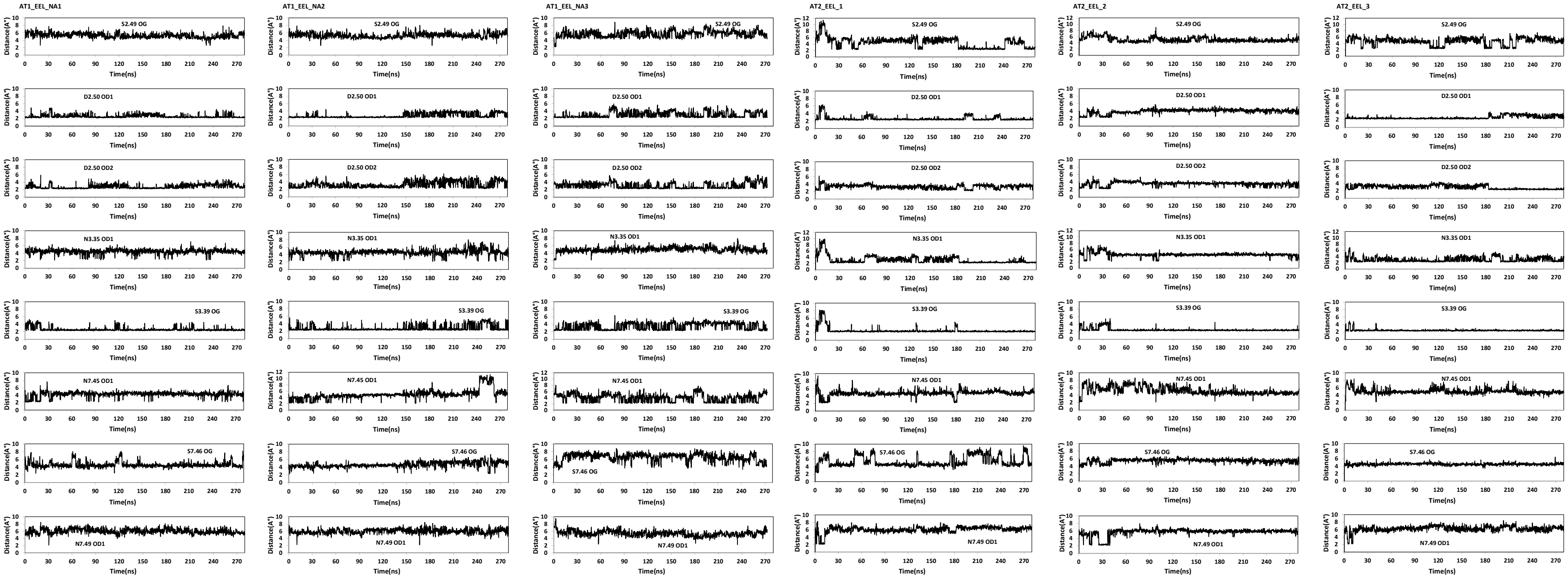
AT1_EEL_NA1
AT1_EEL_NA2
AT1_EEL_NA3
AT2_EEL_1
AT2_EEL_2
AT2_EEL_3
S2.49 OG
D2.50 OD1
D2.50 OD2
N3.35 OD1
S3.39 OG
N7.45 OD1
S7.46 OG
N7.49 OD1
Distance(A°)
Time(ns)

**Fig. S5:** Rotamers of N3.35 in the MD simulations of AT1 and AT2 from human and eel, and the CHIM and N7.46S AT1 mutants.

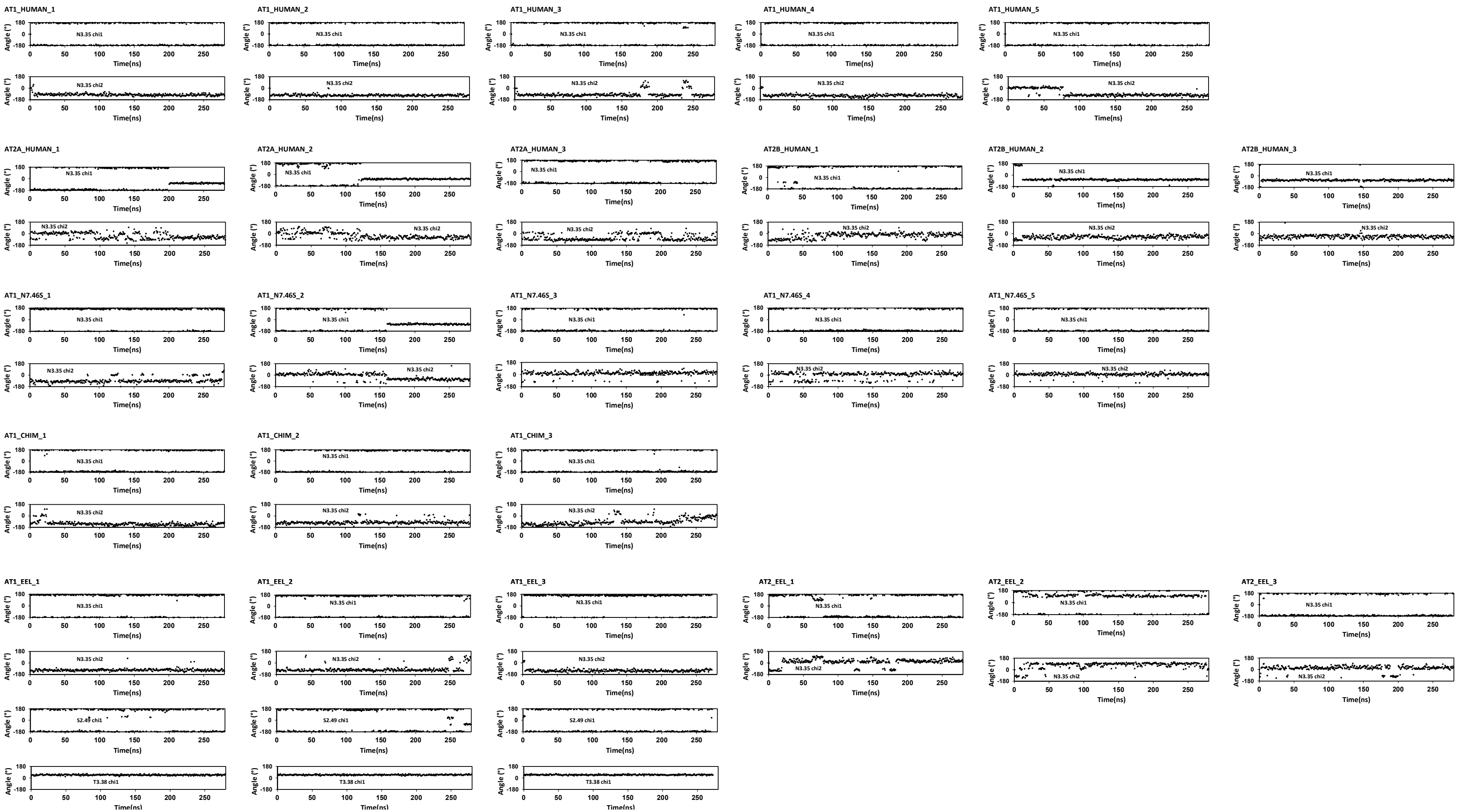
AT1_HUMAN_1
AT1_HUMAN_2
AT1_HUMAN_3
AT1_HUMAN_4
AT1_HUMAN_5
AT2A_HUMAN_1
AT2A_HUMAN_2
AT2A_HUMAN_3
AT2B_HUMAN_1
AT2B_HUMAN_2
AT2B_HUMAN_3
AT1_N7.46S_1
AT1_N7.46S_2
AT1_N7.46S_3
AT1_N7.46S_4
AT1_N7.46S_5
AT1_CHIM_1
AT1_CHIM_2
AT1_CHIM_3
AT1_EEL_1
AT1_EEL_2
AT1_EEL_3
AT2_EEL_1
AT2_EEL_2
AT2_EEL_3
N3.35 chi1
N3.35 chi2
S2.49 chi1
T3.38 chi1
Angle (°)
Time(ns)
